# A Turing-Style Test for In-Silico Antibodies: How Sampling Mode Makes WGAN-GP Beat VAE in the Wet Lab

**DOI:** 10.64898/2026.09.11.750936

**Authors:** Arkadij Kummer, Farbod Mahmoudinobar, Wendi Liu, J. Wade Davis, Eric J. Ma, Sandeep Kumar

**Affiliations:** Computational Science, Research and Early Development, Moderna Inc., 325 Binney Street, Cambridge, MA, 02142, USA; Data Science and Artificial Intelligence (Research), Moderna Inc., 325 Binney Street, Cambridge, MA, 02142, USA; Digital for Research, Moderna Inc., 325 Binney Street, Cambridge, MA, 02142, USA

**Keywords:** Antibody, Deep Learning, Turing Test, Drug Discovery, Biotherapeutics, Artificial Intelligence

## Abstract

Recently, it has become feasible to generate antibodies in silico using AI-based approaches such as deep learning, natural language processing, and diffusion models. This opens the door to computational antibody design as a complement to laboratory-based methods (animal immunization, hybridoma technology, and molecular display) for biologic drug discovery. However, when proposing human antibody sequences, or libraries thereof, it remains essential to determine whether they can be expressed, purified, and biophysically characterized using standardized laboratory assays typically applied early in discovery campaigns. In this work, we devised a Turing imitation game inspired experiment to compare two deep learning methods, Generative Adversarial Networks (GANs) and Variational Autoencoders (VAEs), for generating experimentally viable de novo antibody sequences. A Wasserstein GAN with gradient penalty produced sequences that were successfully validated at a markedly higher rate (92/93; 99%) than those from a VAE (2/40; 5%). These contrasting outcomes can be rationalized by differences in model architecture and sampling strategies. Our findings highlight the promise of computationally driven discovery of antibody-based biotherapeutics.

## 1 Introduction

Antibody-based therapeutics have revolutionized modern medicine, yet their discovery and development remain cost, resource, and time-intensive endeavors with uncertain outcomes. Traditional methods of antibody-drug discovery, including animal immunization and display technologies, often yield candidates requiring extensive downstream engineering to address issues like poor expression, aggregation, or instability. Recent advances in machine learning offer promising alternatives: generative models capable of designing de novo antibody sequences with favorable properties could streamline early-stage development. Towards this goal, we have recently proposed a conceptual roadmap for DAbI (Discovery of Antibodies In-silico) [1] and demonstrated the feasibility of generating highly developable monoclonal antibodies (mAbs) via deep learning [2] via small scale expression, purification and biophysical characterization experiments. However, this initial study was restricted to use of a single deep learning method, namely, Generative Adversarial Neural Networks and did not evaluate other in-silico antibody generation methods.

The field of in-silico antibody generation has witnessed rapid progress in deep generative models for protein design, encompassing architectures such as variational autoencoders (VAEs) [3], generative adversarial networks (GANs) [4], protein language models [5], and diffusion-based frameworks [6]. These models learn complex sequence distributions, enabling the generation of diverse and syntactically valid proteins.

In designing antibodies, approaches differ substantially in how they treat heavy and light chains: a few models are trained on paired heavy-light repertoires (e.g., small subsets of OAS or curated structural-antibody datasets) [7–9], while many large-scale language models and diffusion methods continue to train heavy and light chains independently with chain-type conditioning (rather than explicit pairing) [10, 11].

Despite these advancements, generating antibodies with strong developability profiles remains challenging. Developability encompasses attributes such as protein expression yield, thermal stability, hydrophobicity, aggregation, and poly-specificity. These are critical factors for manufacturability and can also contribute towards clinical success and regulatory approval of biotherapeutic drug candidates. Many studies rely on computational heuristics to assess these properties (e.g. [12]), yet such in-silico scores often correlate poorly with experimental outcomes [13]. When experimental validation is incorporated, it is often focused primarily on binding affinity and expression yield; developability properties are then assessed largely through in-silico predictors, such as TAP scores and surrogate non-specificity classifiers [14].

Recently, high-throughput experimental platforms have been developed to generate large-scale developability datasets in formats suitable for AI/ML training, such as the PROPHET-Ab platform [15], which provides a critical foundation for improving predictive models.

In parallel, an equally important question is whether deep generative models can directly produce antibody sequences with favorable developability profiles. In the spirit of a Turing-style indistinguishability test [16], we ask whether deep generative models can produce antibody sequences that can be expressed, purified, and biophysically characterized such that their developability mimics or exceeds that of natural/marketed antibodies. To probe this, we present a comparative evaluation of two architectures for de novo sequence generation: a VAE and a Wasserstein GAN with gradient penalty (WGAN-GP) [17, 18]. Both models generate paired heavy- and light-chain variable regions (VH and VL, together the Fv), rather than single-domain (VHH) or full-length antibodies; we use VH and VL throughout for these variable regions. Both were trained on 19,096 human IGHV1–IGKV3 paired sequences from the Observed Antibody Space (OAS) database, a pairing common among therapeutically approved antibodies [19].

Our evaluation proceeds in two stages. First, we sample näıve sequences from the VAE and GAN trained on the 19,096 IGHV1–IGKV3 set to characterize fundamental generative behavior: up to ∼1 million sequences per model for the sequence-diversity (rarefaction) analysis, with nearest-neighbor distance to training data and germline annotation accuracy characterized on a ∼500,000-sequence subset. Second, we refine training using a curated subset of 5,480 sequences with high medicine-likeness scores, where medicine-likeness is defined by a composite score from our internal pipeline that integrates calculated sequence-and structure-based descriptors of developability. We then retrain both models on this curated set and sample 2,000 sequences from each for computational profiling. For experimental testing, we selected 133 candidates (93 GAN-generated, 40 VAE-generated) alongside 122 marketed therapeutic antibodies. GAN variants were sampled directly from the retrained model, while VAE variants were generated using latent seeds corresponding to marketed antibodies not included in training, with the goal of probing generalization. This methodological distinction, as shown later in our results, led to differences in medicine-likeness profiles and experimental success rates. Our focus throughout is on intrinsic physicochemical attributes of the generated antibodies; we do not assess target specificity in this work.

## 2 Results

### 2.1 Training deep generative models on human antibodies

We began with a dataset of 19,096 paired antibody sequences from the OAS database. To ensure data quality, sequences were filtered for alignment integrity and correctness, and we focused on the IGHV1–IGKV3 germline family, a pairing that appears frequently among FDA approved antibodies [19]. We refer to this full 19,096-sequence training set as the germline set, distinguishing it from the curated high medicine-likeness subset (n=5,480) used for retraining.

On this dataset, we trained two deep learning models with contrasting architectures. The Variational Autoencoder (VAE) compresses antibody sequences into a lower-dimensional latent space and then decodes new variants by sampling around these latent points. This structure tends to keep generated sequences close to training data but can be sensitive to how sampling is performed. The Wasserstein Generative Adversarial Network with Gradient Penalty (WGAN-GP) instead generates antibodies directly from random noise, with a generator network proposing new sequences and a discriminator network teaching it to mimic natural antibodies more closely. This adversarial setup often produces diverse and realistic samples but can be harder to train stably (see Methods for further details).

The fundamental difference in how the models generate sequences is important to highlight. GAN sequences are sampled directly from random noise, meaning that every sample is independent and unconditional. VAEs, in contrast, require a seed sequence: they encode an antibody into a latent representation and then decode nearby variants. This seed-dependence has practical consequences for both diversity and experimental outcomes, as shown below.

Both models trained successfully without signs of instability or overfitting. The VAE steadily improved its reconstruction accuracy while maintaining a well-regularized latent space. The GAN reached a stable balance between generator and discriminator, avoiding the mode collapse sometimes seen in adversarial learning (Supplementary Figure 1).

### 2.2 GANs produce broader and more diverse antibody libraries

To test how well each model could explore antibody sequence space, we sampled one million naive VH/VL pairs from each. We first measured internal diversity, the fraction of samples distinct from one another: GANs produced nearly all unique pairs (99.99%), while only ∼41% of VAE samples were distinct, the rest being repeats. This saturation was especially pronounced in light chains, where many VAE samples were repeated (Fig. 1, panels A-C).

**Fig. 1.**
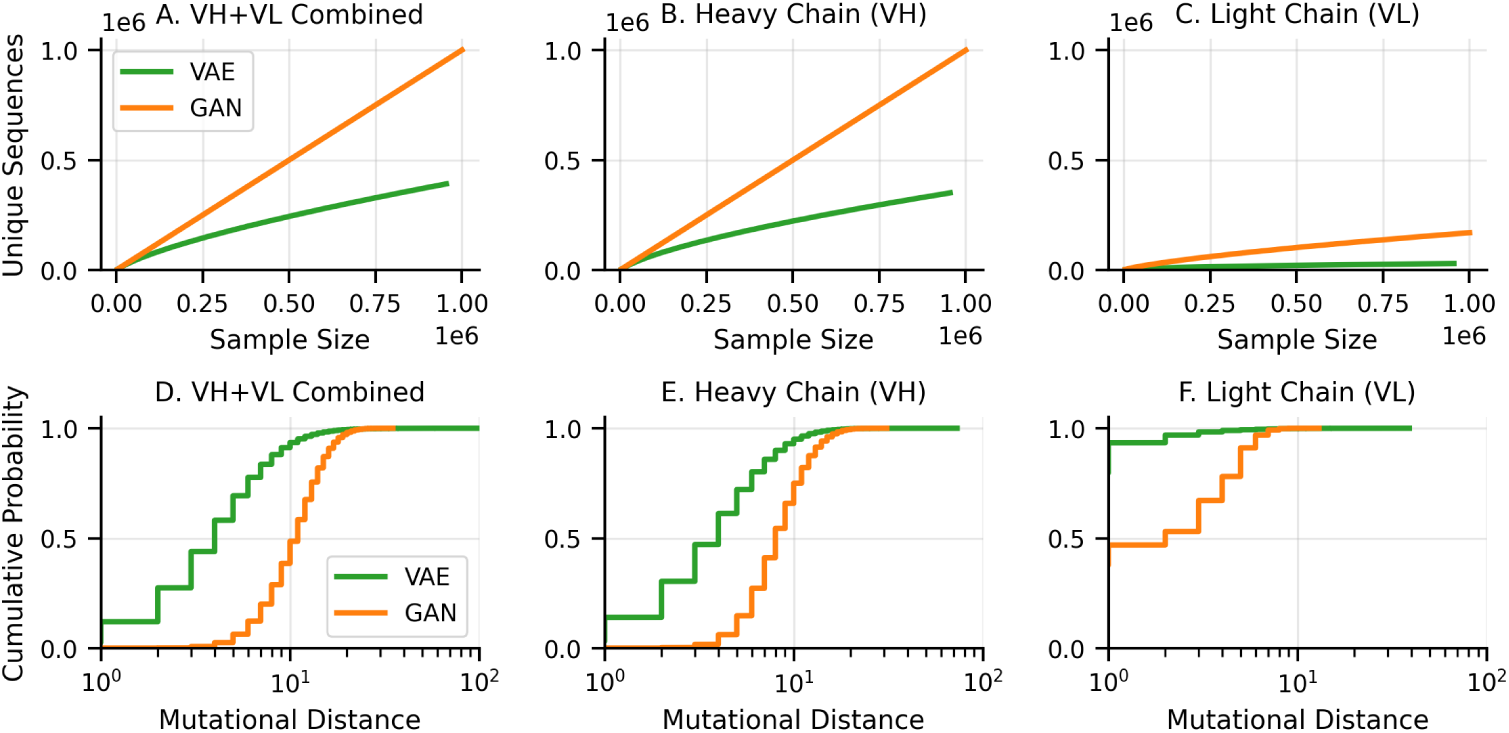
Model Performance Comparison. Panels A-C show rarefaction curves of cumulative unique sequences versus sample size for ∼1M samples/model across combined VH+VL, VH only, and VL only. GAN (orange) reaches 999,940 unique pairs (99.99%) while VAE (green) reaches 391,433 (41.0%), indicating near-linear diversity growth for GAN and early saturation for VAE. Panels D-F show empirical CDFs of mutational distance (Hamming distance to nearest training sequence) for the same sequence types. VAE sequences cluster closer to training data (mean ∼5 mutations) while GAN sequences explore further (mean ∼10 mutations), both within ranges consistent with natural somatic hypermutation.

This difference reflects the sampling strategies. Because GANs sample unconditionally from noise, they naturally cover sequence space more broadly. VAEs, by contrast, are constrained by the seeds used for decoding: sampling around the same or similar seeds repeatedly leads to concentrated, less diverse libraries. This explains the early saturation observed in VAE rarefaction curves compared to the near-linear diversity growth of GANs.

### 2.3 Both models preserve natural antibody features

Despite differences in diversity, both models generated antibodies that preserved natural architecture and sequence features. We used the ANARCI python package [20], which assigns standard positions to antibody residues, to check for structural validity. We chose to use the Martin numbering scheme for this. Every generated sequence had correct domain boundaries and plausible lengths (Fig. 2, panel A).

**Fig. 2.**
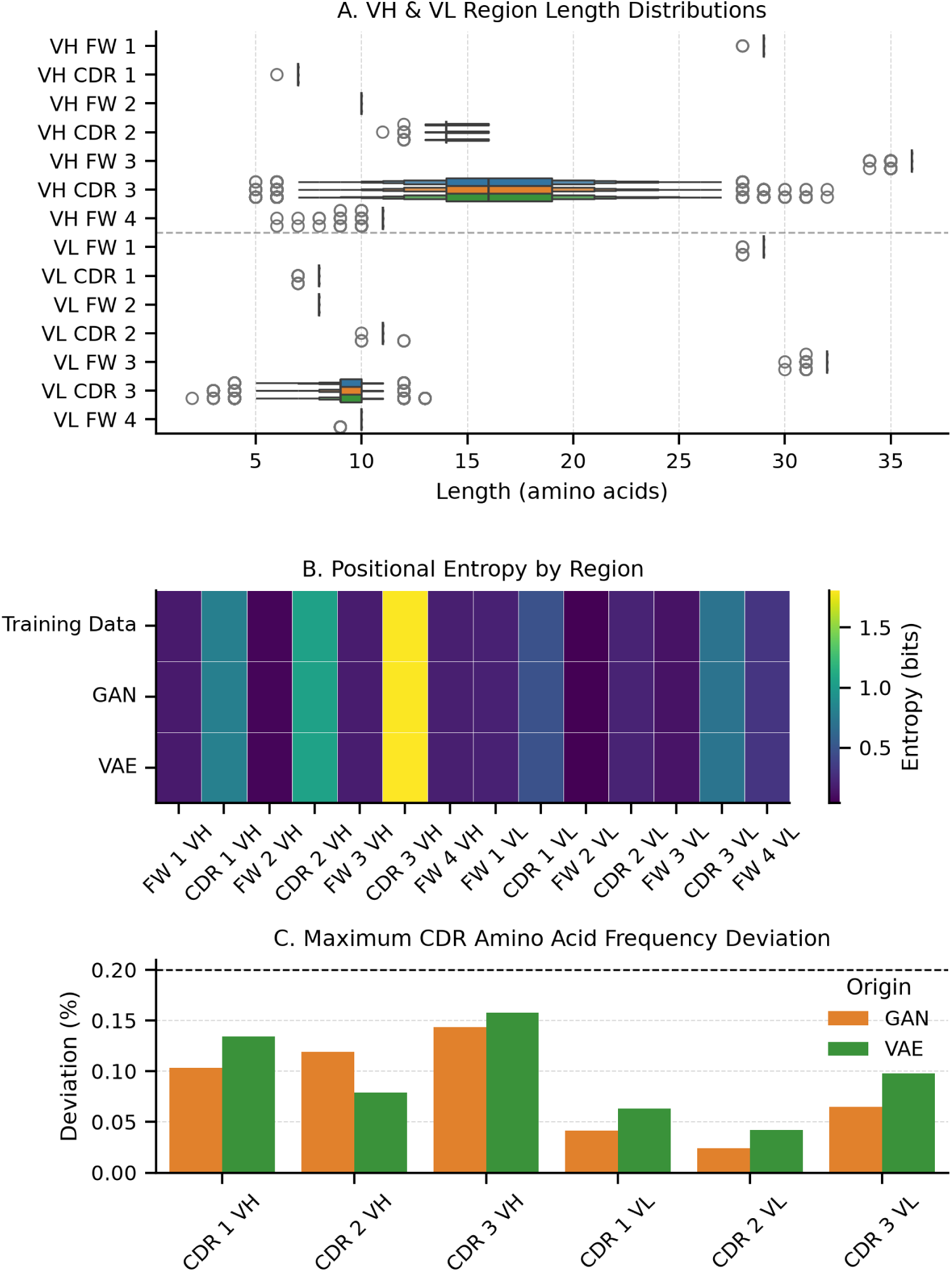
Sequence Quality Validation. Panel A shows VH and VL region length distributions under Martin numbering for training data, VAE, and GAN samples. Boxplots show median, IQR, and outliers for FR1-FR4 and CDR1-CDR3 regions with a dashed line separating heavy and light chains. Both models closely match training profiles with narrow frameworks and broader CDRs, especially CDR3. Panel B shows average positional entropy by region, confirming high CDR variability and conserved framework regions across training data and generated samples. Panel C shows the maximum CDR amino acid frequency deviation from the t6raining distribution for each model; all deviations remain below 0.2%.

When projected into two dimensions using Uniform Manifold Approximation and Projection (UMAP), both GAN and VAE samples formed distributions that closely resembled the original training set, indicating that both models had learned the broad global features of antibody repertoires.

Closer inspection confirmed that biologically important patterns were preserved. The characteristic high variability of the complementarity-determining regions (CDRs), especially CDR3, was maintained, while framework regions remained conserved (Fig. 2, panel B). Amino acid usage in CDRs nearly matched natural frequencies, with deviations of less than 0.2% (Fig. 2, panel C). Germline identity was preserved exactly, with both models retaining the IGHV1–IGKV3 family constraints and reproducing natural J-gene usage (Supplementary Figure 2).

### 2.4 Sampling behavior affects mutational distances

Novelty is a distinct question from internal diversity: for a ∼500,000-sequence sample from each model, what fraction of the distinct sequences is absent from the training set? By this measure both models are highly novel, GAN 100% and VAE 98.1% (Supplementary Figure 3). Thus the VAE’s limitation is redundancy rather than memorization: it revisits the same sequences often (only ∼41% distinct in the diversity analysis above), yet the distinct sequences it does produce are almost all new relative to training. Heavy chains were more novel than light chains in both models (Supplementary Figure 3).

We next compared how far generated antibodies were from their nearest training neighbors. Using normalized Hamming distance over the aligned sequences, we found that VAE sequences were closer to training data (mean 4–5 mutations) than GAN sequences (mean ∼10 mutations) (Fig. 1, panels D-F). Both ranges are consistent with natural somatic hypermutation in B-cell repertoires [21].

However, the VAE occasionally produced extreme outliers with more than 50 mutations. Interestingly, these outliers arose not only when we increased the latent log-variance (which controls how far the decoder samples from the seed embedding), but also under the standard setting (log-var multiplier = 1.0). This suggests that when seeds lie at the edge of the learned distribution, even moderate perturbations can push decoding into unsupported regions, producing degenerate outputs (Supplementary Figure 4). GANs, by contrast, showed broader but more uniformly distributed mutational distances, consistent with their unconditional sampling. This indicates that GANs naturally explore further from the training set without venturing into implausible territory.

### 2.5 Training on medicine-like antibodies enriches libraries

We then retrained both models on a curated subset of 5,480 sequences with high medicine-likeness, defined using nine descriptors of developability such as aggregation propensity, charge distribution, and hydrophobic balance. These descriptors were benchmarked against 122 variable regions from marketed antibodies, and scores were normalized to allow comparison. The medicine-likeness scoring pipeline is described in detail by Mahmoudinobar et al. [22].

From these retrained models, we generated 2,000 sequences each for in-silico analysis. GAN sequences were sampled unconditionally, while VAE samples were seeded with training sequences to ensure variants remained on-manifold. Both models produced libraries with higher average medicine-likeness than the germline set (mean scores: VAE 1.57, GAN 1.25, vs. 0.30 for the germline set). Both also generated individual antibodies exceeding the maximum scores of the high medicine-likeness subset, showing their ability to generate novel, highly medicine-like variants.

Encouraged by these results, we again sampled antibodies from the retrained models to send for experimental characterization. It is important to note, however, that the experimental subsets were sampled differently. GAN variants were drawn in the same unconditional manner. VAE variants were instead seeded with marketed antibodies not included in training. These “out-of-distribution” seeds led to weaker, often degenerate sequences with lower medicine-likeness scores, in contrast to the strong in-silico results obtained when VAEs were seeded with training data. Those in-silico samples (VAE n=2,012; GAN n=2,000) are the distributions shown in Fig. 3; the weaker experimentally tested subset is characterized in the following section.

**Fig. 3.**
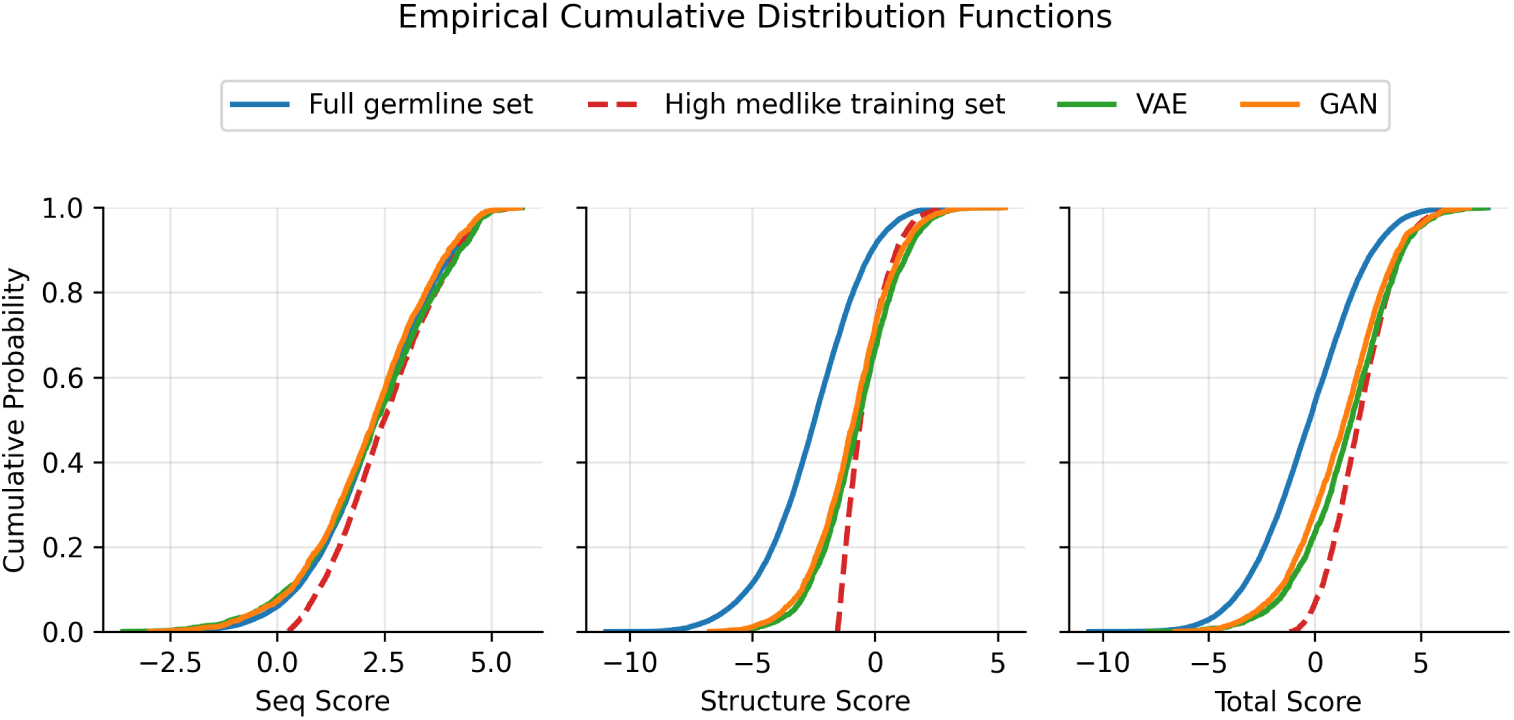
Medicine-likeness Score Distributions. Empirical cumulative distribution functions (ECDFs) of sequence, structure, and total medicine-likeness scores. Curves show the full germline set (n=19,096), the high medicine-likeness training subset (n=5,480), and the model samples (VAE n=2,012; GAN n=2,000). Both generators show right-tail extensions beyond the subset maximum, with total-score means placing the sampled generators between germline and subset levels.

### 2.6 Laboratory testing shows robust GAN success, weak VAE outcomes

To validate experimental tractability, we selected 133 generated antibodies (93 GAN, 40 VAE) alongside 122 marketed antibodies. For each, expression as IgG1 in Chinese Hamster Ovary (CHO) cells, purification, and evaluation in nine standardized assays covering expression yield, aggregation, thermal stability, hydrophobicity, self-association, and non-specific binding were attempted; not all candidates expressed successfully (see below). For more details on the experimental evaluation, see Liu et al (in preparation).

Although detailed results are reported separately, the overall trend was clear. GAN-derived antibodies expressed successfully and produced biophysically tractable proteins across assays, with many GAN variants comparable to marketed antibodies in the measured developability assays. The experimentally tested VAE candidates performed poorly, with most not expressing. These outcomes should be interpreted in light of the sampling differences described above: the experimental VAE subset was seeded from out-of-training marketed antibodies, whereas the in-silico VAE library seeded from training sequences showed considerably stronger medicine-likeness profiles (see Fig. 4, panel B). The comparison therefore reflects the sampling protocols applied to each model rather than an architecture-only contrast.

**Fig. 4.**
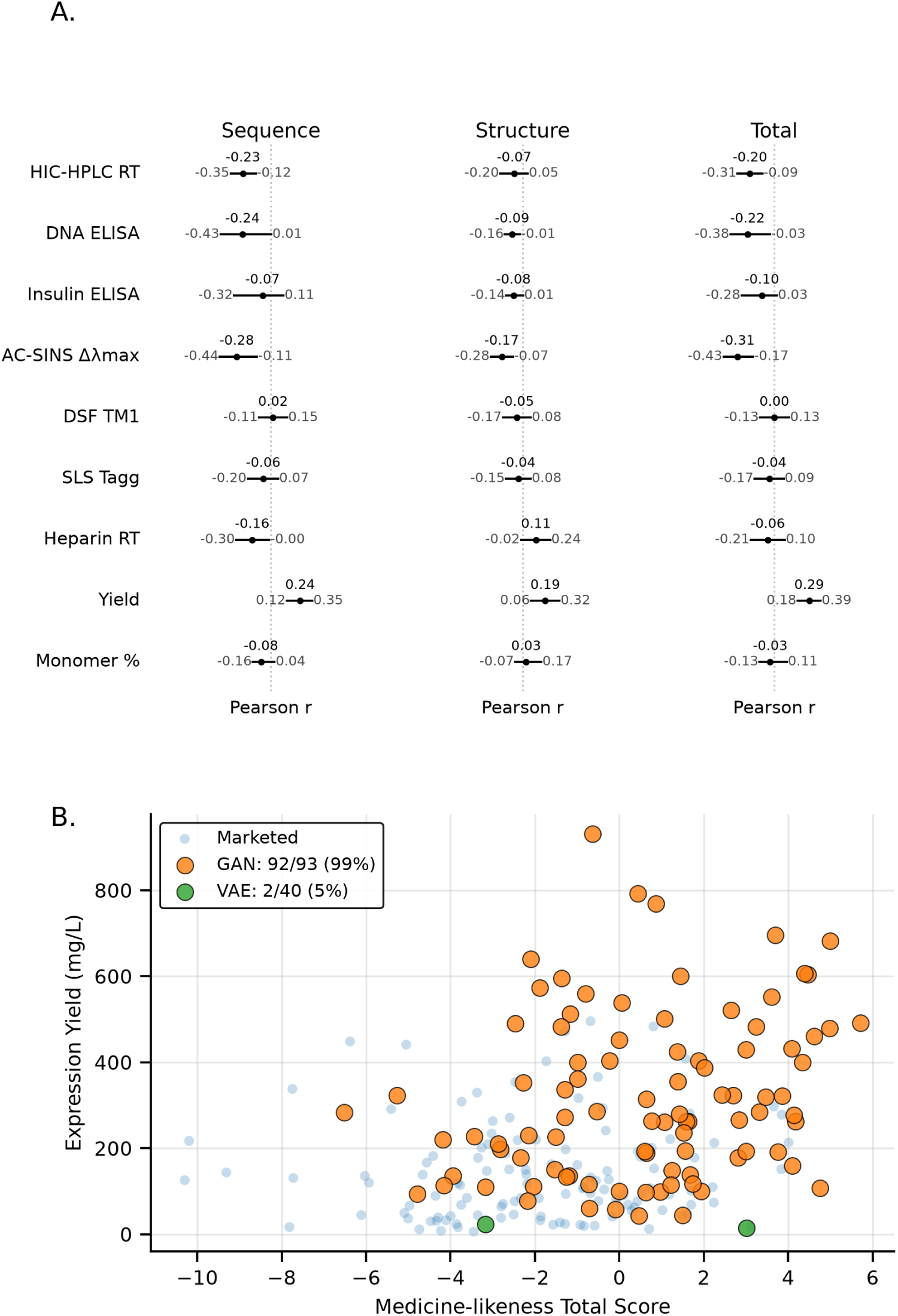
Experimental Results. (A) Pearson correlation coefficients with 95% bootstrap confidence intervals (10,000 iterations) between computed m1e0dicine-likeness descriptors and experimental measurements for 226 antibodies (90 GAN-generated, 109 marketed, and 27 additional non-VAE reference antibodies comprising engineered variants, clinical-stage benchmarks, withdrawn therapeutics, and the NISTmAb standard; VAE candidates excluded due to poor experimental performance). Correlations are generally weak (mean |r| = 0.13), with the strongest being 0.31 between total score and AC-SINS Δ*λ*max. (B) Expression yield versus medicine-likeness total score. GAN-generated antibodies (orange, n=93) show 99% expression success (92/93), while VAE-generated antibodies (green, n=40) show only 5% success (2/40). Marketed therapeutic antibodies (blue, n=122) serve as reference.

To assess whether medicine-likeness descriptors correlate with experimental outcomes, we computed Pearson correlation coefficients with bootstrap confidence intervals between simulated properties (sequence score, structure score, total score) and all nine experimental assays for the 226 successfully characterized non-VAE antibodies (90 GAN-generated, 109 marketed, and 27 additional reference antibodies; VAE candidates excluded). Fig. 4 (panel A) shows the complete correlation analysis. The correlations were generally weak across all property pairs. The strongest observed correlation was 0.31 [0.43, 0.17] between total score and AC-SINS Δ*λ*_max_, while most correlations had magnitudes below 0.25 with confidence intervals spanning near-zero values. Expression yield showed positive correlations with all three scores (range: 0.19– 0.29) but with wide confidence intervals reflecting substantial variability. These weak correlations indicate limited predictive power of computational descriptors for experimental developability outcomes, consistent with recent findings showing that in-silico physicochemical metrics often correlate poorly with measured antibody developability [13].

### 2.7 Predictive modeling reveals limited correlation between in-silico descriptors and lab results

Finally, we tested whether the biophysical descriptors from our medicine-likeness pipeline could predict experimental assay outcomes. Using the 122 marketed antibodies and 93 GAN-generated antibodies, we trained machine learning models to predict individual assay values based on the in-silico descriptors. We decided to exclude the VAE generated antibodies here due to their overall degeneracies and weak performance. In plain terms, we asked whether quick, computation-only scores could help flag sequences that are more likely to behave well in the lab before running the assays. In detail, we computed an array of numerical descriptors capturing medicine-likeness, physicochemical properties, liabilities, and immunogenicity risks. This feature array was used as input to predictive models for a five-assay subset of the broader experimental panel: Hydrophobic Interaction Chromatography (HIC) retention time, Affinity-Capture Self-Interaction Nanoparticle Spectroscopy (AC-SINS) peak wavelength shift (Δ*λ*_max_), Differential Scanning Fluorimetry (DSF) first melting temperature (*T_M_*_1_), Static Light Scattering (SLS) aggregation onset temperature (*T_agg_*), and expression/purification yield. In other words, each target variable in this predictive benchmark was modeled in turn from the full descriptor set.

To guard against memorization of near-identical sequences, we used sequence-similarity–blocked cross-validation (see Methods), in which the similarity threshold sets how alike a test sequence may be to any training sequence. A lower threshold (0.8) forces test sequences to be more dissimilar from training and thus poses a harder generalization task than a higher one (1.0). Under this scheme, the individual descriptors used to construct the medicine-likeness profile showed weak regression signal but modest binary-classification signal (Fig. 5).

**Fig. 5.**
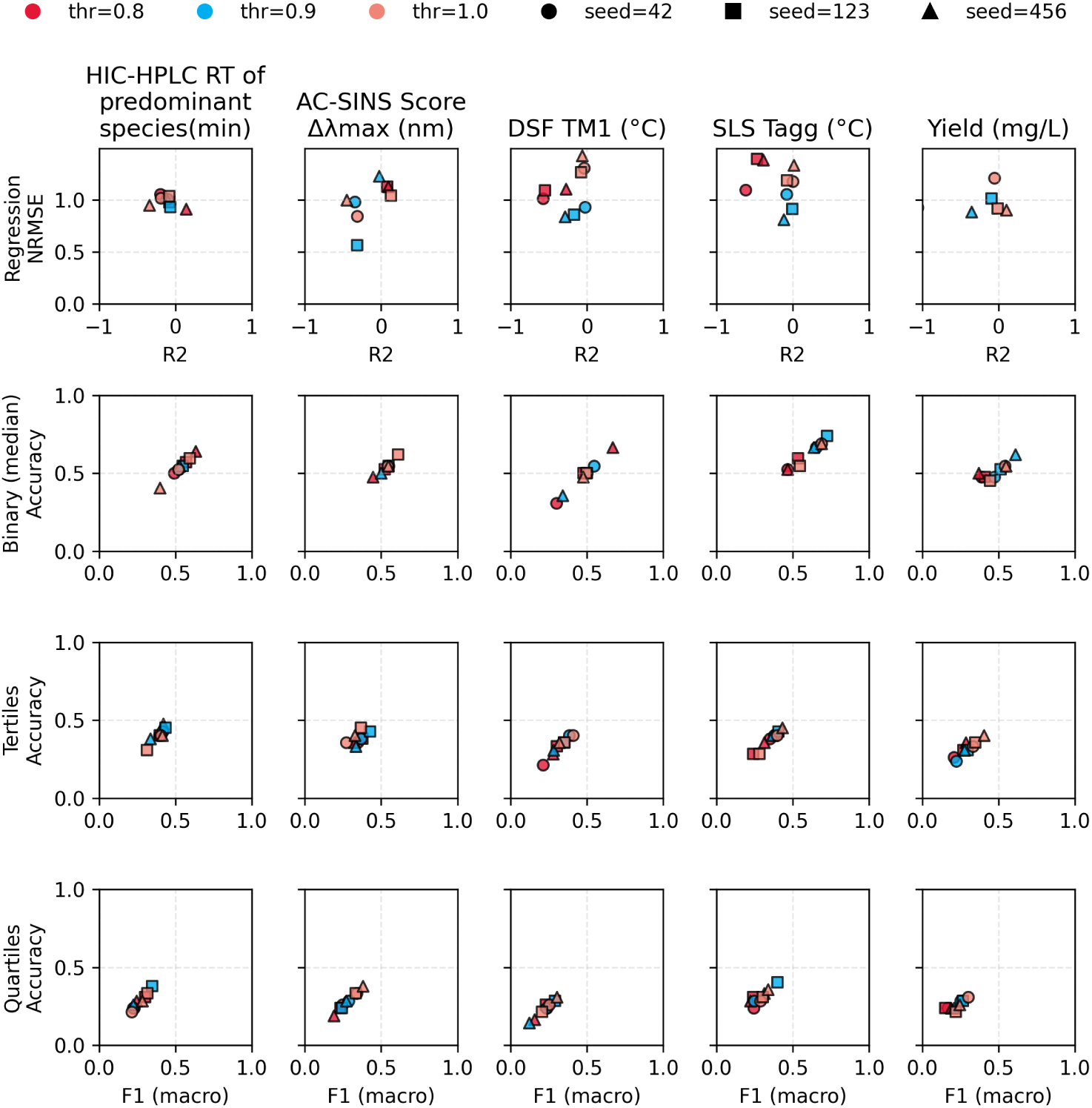
Predictive Model Performance. Predictive performance of medicine-likeness descriptors under sequence-similarity–blocked cross-validation across five assays (HIC-HPLC RT, AC-SINS, DSF TM1, SLS Tagg, Yield). Colors encode similarity thresholds (0.8, 0.9, 1.0); marker shapes encode three random seeds (42, 123, 456). The top row shows Ridge regression *R*^2^ versus NRMSE, and the lower rows show Random Forest classification macro-F1 versus accuracy for binary, tertile, and quartile outcome bins. Performance clusters near chance baselines, indicating limited predictive utility of computational descriptors for experimental outcomes.

In Ridge regression, runs cluster near *R*^2^ *≈* 0 with errors comparable to a mean predictor; the best single split reached *R*^2^ = 0.234 for AC-SINS at the 0.9 threshold. Typical error ranges were large across assays (for example, HIC-HPLC RT RMSE 2.43–2.67 min, DSF TM1 4.40–4.90 °C, SLS Tagg 6.99–7.67 °C, Yield 167–175 mg/L). Predicted-versus-actual plots show regression to the mean and underestimation at higher observed values (Supplementary Figure 6).

In contrast, Random-Forest binary (median) classifiers captured some signal. Here, “median” means we binned each assay’s y-values into two groups and predicted whether a sequence fell below or above that assay’s median value. Likewise, “tertiles” and “quartiles” refer to splitting the y-values into three or four ordered bins for multiclass classification. Under this setup, median prediction macro-F1 and accuracy typically fell in the 0.54–0.66 and 0.59–0.66 ranges for HIC-HPLC RT, AC-SINS, SLS Tagg, and Yield, with a best case of AC-SINS macro-F1 = 0.714 and accuracy = 0.714 at threshold 1.0 (Fig. 5); DSF TM1 remained closer to chance (macro-F1 about 0.46– 0.53). Performance declined for finer granularity (tertiles and quartiles), and most classification errors occurred between adjacent bins across thresholds (Supplementary Figure 5).

## 3 Discussion

This study was designed to address two central questions. First, in the spirit of a Turing-style test, which generative model is more likely to produce antibody sequences that survive the practical hurdles of laboratory expression and characterization? Second, given that we profiled antibodies both computationally and experimentally, can in-silico descriptors be reliably used to predict experimental outcomes?

To answer these questions, we compared two widely used generative architectures, WGAN-GP and VAE, trained on antigen-agnostic antibody libraries from the IGHV1– IGKV3 germline family.

We chose sequence-only GANs and VAEs as a pragmatic fit for our goals: generating very large, antigen-agnostic libraries within a single germline family and analyzing distributional properties (novelty, diversity, region lengths, germline fidelity) at scale. They are straightforward to train on tens of thousands of sequences, inexpensive to sample, and let us run controlled comparisons (unconstrained vs. implicitly curated “medicine-like” training) without added supervision. Current antibody LLMs and diffusion models offer finer-grained control (conditional or structure-aware design) but carry heavier requirements, such as larger corpora, more compute, and structure or paired-chain conditioning, that were not needed for our core questions. GANs and VAEs thus gave us fast, reproducible baselines matched to this work’s scope; LLMs and diffusion remain promising for future epitope- or structure-aware design.

This controlled setting allowed us to assess core properties such as sequence diversity, novelty, and germline fidelity at scale. At first glance, GAN-generated antibodies appeared more tractable in experimental evaluation, while VAE-derived antibodies underperformed. However, this difference should not be viewed as a simple architectural comparison; it is tightly linked to how each model is sampled in practice. GANs generate antibodies unconditionally: every sequence is created directly from random Gaussian noise, which makes the full repertoire equally accessible and provides a robust, reproducible way to explore sequence space. VAEs, by contrast, are inherently seed-dependent. A seed sequence is encoded into a latent vector, and new antibodies are produced by decoding perturbations around that point. The effectiveness of this process is therefore highly sensitive to which seeds are chosen and how far sampling is allowed to wander in latent space.

This methodological distinction shaped both our in-silico and experimental outcomes. In large-scale computational profiling, where VAE seeds were drawn from the training set, the resulting libraries were competitive: they preserved germline constraints, achieved reasonable novelty, and reached medicine-likeness scores comparable to GAN libraries. But for the experimental evaluation, VAE seeds were taken from marketed antibodies not present in the training set. This choice was meant to test generalization but had unintended consequences. It pushed the VAE decoder into lower-support regions of latent space, where perturbations more easily yield degenerate or implausible sequences. Hence, the experimentally tested subset of VAE sequences underperformed relative to GAN sampled sequences. By contrast, GANs, being seed-free, avoided these pitfalls and consistently produced diverse, on-manifold candidates that were more straightforward to express and characterize, with developability characteristics comparable to those of approved antibodies in the measured assays.

This latent-space challenge persists even when generating antibodies within a single germline family, as performed in our work. Recent work shows that antibody generative models can memorize germline residues while underrepresenting the non-germline variation that distinguishes functional antibodies, a specific risk when training is confined to one germline family [23]. Unlike enzyme engineering, where the latent space around a single functional scaffold is often dense with viable variants that tolerate many local moves during optimization (especially away from the active site residues) [3, 24], the antibody landscape is considerably more brittle: small sequence changes can perturb paratope geometry, surface patches, or packing features that matter for stability and aggregation. As a result, VAE sampling without strong guidance readily drifts into low-support or invalid regions, whereas adversarial training in WGAN-GP empirically favors samples that better respect the full training distribution. Prior enzyme studies illustrate how scaffold-centered latent modeling can successfully yield many functional variants (for example, VAE-guided libraries for ornithine transcarbamylase and other enzymes), underscoring how antibodies present a tougher generative target despite conservation of the *β*-barrel fold even among sequence divergent molecules [3, 25].

The comprehensive comparison between VAE and GAN architectures revealed fundamental differences in their practical utility for antibody library generation. Our rarefaction analysis demonstrated that the GAN achieved near-perfect sequence diversity compared to the VAE’s substantially lower uniqueness rate, with particularly striking differences in light chain generation. While the GAN enables straightforward scaling to millions of unique candidates, the VAE requires sophisticated sampling strategies to overcome mode-collapse-like redundancy and to maintain validity. In practice, this means the GAN is better suited for broad, nonredundant exploration, whereas the VAE benefits from deliberate, seed-centered or density-aware strategies and, where appropriate, conditional formulations that encode task-relevant priors. Prior antibody-focused VAE work has highlighted that remedies like conditioning and density-aware priors can materially improve sample quality, though they introduce additional modeling and operational complexity [26].

Notably, GAN-generated antibodies showed experimental properties on par with or better than marketed therapeutics across multiple biophysical assays. However, because our sequence-aware prediction effort using medicine-likeness descriptors as inputs to predict experimental readouts showed only modest signal, we avoid attributing the GAN’s favorable wet-lab behavior solely to the implicitly curated training subset. More plausible contributors include preservation of natural antibody constraints learned from the germline family, the IGHV1-IGKV3 framework context, and inductive biases of adversarial training that discourage off-manifold samples. The weak descriptor-to-assay correlations reported above caution against inferring causality from score improvements, and dedicated machine-learning developability predictors report similarly mixed, endpoint-dependent accuracy: PROPERMAB predicts hydrophobic-interaction retention time moderately well (Pearson *r ≈* 0.71) while predicting high-concentration viscosity only weakly (*r ≈* 0.35) [27]; structure-based tools built on AlphaFold2 models [28] and multi-encoder or text-augmented frameworks [29, 30] improve some endpoints while indicating that properties such as expression yield and thermal stability depend on paired heavy–light context. Coupling such predictors to guided generation has been proposed to enrich designed libraries for developability [31], though experimental confirmation across a full assay panel remains the limiting step.

To keep our performance estimates realistic, we split the data so that closely related sequences never appeared in both the training and test sets. With this kind of sequence-similarity aware cross-validation, apparent predictability often drops compared with a conventional random split, which can give overly optimistic results. This concern is common in biomolecular machine learning: if near-duplicate or homologous sequences show up in both sets, similarity can be mistaken for learnable signal and inflate reported metrics [32]. In our case, because the true predictive signal was already weak, we can only make a modest claim about how much our choice of cutoffs for binning the outcomes (for example, above or below the median) affected the results. Even so, we recommend sequence-aware splitting as a safeguard against homology leakage and as a more realistic test of generalization.

These results arrive amid rapid progress in deep generative protein design. Antibody-specific language models trained on large repertoires, including AbLang [33], AntiBERTy [34], and the paired-chain IgBert and IgT5 models [5], together with general protein language models such as ESM-2 [35] and ProGen2 [36], provide realistic sequence priors and can aid mutation suggestion or gap-filling. Paired-chain generative models now produce antigen-specific heavy–light pairs with experimental binding validation, as demonstrated by MAGE [8]. Diffusion-based and inverse-folding methods for antibody sequence–structure design, including DiffAb [37], AbDiffuser [38], IgDiff [39], ProteinMPNN [40], and AntiFold [41], have expanded control over loops, paratope geometry, and chain pairing, and recent foundation models combine diffusion with language modeling over paired chains [9]. De novo design platforms now report experimental results at scale: Chai-2 reports a 16% binding hit rate across 52 targets [42], and JAM achieves therapeutic-grade yield, monomericity, and low polyreactivity for de novo VHHs, though it is VHH/nanobody-oriented and epitope-specific and its designs remain at humanized-level germline identity [43]. Most recently, Latent-X2 has begun to combine all three validation axes in one system, reporting zero-shot antigen-specific binders (VHH and scFv) with developability profiles matching approved therapeutic controls and, for the first time for a de novo AI-generated antibody, ex vivo immunogenicity assessment in human donor panels [44]. Benchmarking efforts such as AbBiBench [45] and the structure-focused CHIMERA-Bench [46] have begun to compare families of language and diffusion models specifically for antibody design, though most evaluations still rely heavily on in silico metrics and limited experimental readouts.

It is important to situate the present work correctly. The de novo binder platforms above (Chai-2, MAGE, JAM, Latent-X2) are antigen-specific: they design against a defined target and epitope, and are increasingly validated along binding, developability, and immunogenicity axes at once. Our study asks a complementary question: the intrinsic, antigen-agnostic developability and wet-lab tractability of paired variable regions (VH/VL) generated within a single germline family and expressed as full-length IgG1, without any binding or functional selection. We therefore do not treat these binder-discovery platforms as direct comparators, and in-silico-focused systems should not be placed in the same evidentiary class as wet-lab-validated ones. Rather, the trajectory they define, coupling generation to multi-axis experimental validation, is the direction in which antigen-agnostic developability profiling such as ours can feed as an upstream, target-independent filter.

As a cautionary counterpoint, recent work in enzyme design systematically assessed whether computational metrics actually predict experimental behavior at scale. [47] evaluated a broad panel of in silico scores against the laboratory performance of more than 500 natural and AI-generated enzymes, showing that only through extensive expression and testing could useful filters be identified and refined; a research briefing summarized how a composite filter (COMPSS) improved success rates but still required large-scale experiments to establish reliability. Other recent work by [14] has begun to operationalize this integration through “lab-in-the-loop” systems that orchestrate generative models, multi-task property predictors, active-learning–based ranking/selection, and iterative in vitro measurements in a semi-autonomous loop, enabling end-to-end, multi-property optimization under real experimental feedback. These frameworks exemplify how rapid experimental iteration can directly inform model updates and selection criteria, narrowing the gap between in-silico rankings and manufacturable, well-behaved antibodies. The lesson generalizes: comprehensive experimental campaigns remain essential to calibrate and stress-test computational metrics before relying on them in design loops.

Taken together, our results support a cautious but optimistic path forward. Generative models can indeed create antibody libraries enriched for developability, but success depends as much on sampling strategy and validation design as on the underlying architecture. Experimental evaluation remains essential but must be interpreted considering its variability and limitations. The most promising way ahead is not to rely solely on computation or experiment, but to integrate them: use broad, unconstrained generative training to ensure diversity; refine with curated, property-enriched data; apply sequence-aware validation to avoid homology leakage; and pair modern generative models with iterative, high-throughput experimental characterization in a lab-in-the-loop workflow. This integrative approach, rather than treating computational scores or wet-lab assays as definitive in isolation, will most reliably yield antibody libraries that are both diverse and genuinely developable.

## 4 Methods

### 4.1 Model architectures

Our Variational Autoencoder (VAE) followed the design of [3], adapted for antibody sequences. Prior to training, each scFv or VHH was either Martin-aligned using ANARCI or right-padded with gap symbols so that all examples shared a common length L. Sequences were one-hot encoded over an alphabet of 20 amino acids plus a gap character, producing binary tensors of shape (A, L) that were flattened before entering the network. The encoder comprised four linear layers with ReLU activations (default widths 256 128 96 64), ending in two 64-unit heads that output the mean (*µ*) and log-variance (log *σ*^2^) of the approximate posterior *q*(*z|x*). A latent vector *z* was drawn via the reparameterization trick and decoded through a mirror-image network (64 96 128 256) reshaped back to (A, L) and converted to categorical probabilities with a SoftMax. Optimization used Adam with AMSGrad, a gradient clip of 10*^−^*^6^, and a loss combining negative log-likelihood with an annealed KL-divergence term whose weight followed a sigmoidal schedule. Early stopping halted training after five epochs without validation improvement. At inference, the trained encoder produced *µ* and *σ*^2^ for any seed sequence; by scaling *σ*^2^ with user-chosen factors (0.1, 1, 1.5, 2), we tuned the exploration radius of the latent space before decoding to gapped strings or to ungapped heavy/light chains.

Our Wasserstein GAN with Gradient Penalty (WGAN-GP) was adapted from [2] with sequence-specific handling implemented in PyTorch Lightning’s manual-optimization loop. Sequences were represented as 4-D tensors (batch, alphabet, length, 1) after alignment (ANARCI or right-padding). To ensure compatibility with stride-2 transposed convolutions, both the alphabet size A and sequence length L were padded to the nearest multiple of 16. The generator accepted 128-dimensional Gaussian noise vectors, mapped them through a linear layer into initial feature maps, and progressively up-sampled through four ConvTranspose2d layers with kernels (4Ö4, 4Ö4, 2Ö2, 2Ö2), BatchNorm, and ReLU activations. A cropping layer corrected overshoot from the stride-2 kernels, and a final Sigmoid produced grey-scale logits decoded by argmax back to amino acid symbols, with a gap channel preserved for alignment. The discriminator mirrored this architecture in reverse, employing four convolutional layers with alternating 2Ö2 and 4Ö4 kernels, LeakyReLU activations (*α* = 0.2), and progressive channel expansion (1 C 2C 4C 8C) before a final linear unit estimated the Wasserstein distance. Training followed the WGAN-GP procedure with *λ* = 10 gradient penalty applied to real–fake interpolants, a 5:1 critic-to-generator update ratio, and dynamic padding to maintain consistent tensor shapes. During training, all loss curves and Wasserstein distance metrics were logged for stability assessment.

### 4.2 Model sampling differences

#### VAE Sampling (Seed-Based)

The VAE requires seed sequences from the training data to anchor sampling in the latent space. For each seed sequence, the encoder generates a mean (*µ*) and log-variance (log *σ*^2^) defining a Gaussian distribution in latent space. New sequences are sampled by: (1) encoding the seed sequence to obtain *µ* and log *σ*^2^, (2) applying log-variance multipliers (e.g. 0.1, 1.0, 2.0) to control sampling radius, and (3) drawing samples from the modified Gaussian distribution. This approach inherently limits diversity since all generated sequences are anchored to existing training examples, and the number of unique seeds constrains the total achievable diversity.

#### GAN Sampling (De Novo)

The GAN generates sequences through direct sampling from the learned data distribution without requiring seed sequences. For each desired sample, a random noise vector z is drawn from a standard normal distribution (z ∼ N(0, I)) in the latent space and passed through the generator network. This approach enables diversity scaling that is only constrained by the theoretical sequence space (20^sequence length for natural amino acids) and the generator’s capacity to map latent vectors to valid antibody sequences.

While the VAE can generate multiple diverse sequences from a single seed by varying log-variance multipliers and sampling multiple times from each seed’s latent distribution, the total achievable diversity remains fundamentally bounded by the number of available seed sequences. Even with extensive sampling around each seed (hundreds or thousands of variants per seed), the VAE’s diversity ceiling is determined by how many distinct training sequences can serve as anchoring points, whereas the GAN’s diversity grows with sampling effort without an explicit seed-count constraint. This advantage is not unbounded, however: GAN output diversity is ultimately limited by finite generator capacity, incomplete mode coverage of the training distribution, and the resolution of the sequence decoding step, so continued sampling yields diminishing returns rather than truly unlimited novelty.

### 4.3 Medicine-likeness scoring

The medicine-likeness profile used to curate our training datasets is described in full by Mahmoudinobar et al. [22]; we summarize the essential elements here. Medicine-likeness integrates nine non-redundant descriptors of developability, benchmarked against the reference distribution of 122 unique variable regions (Fvs) drawn from 117 FDA-approved therapeutic antibodies (as of February 2024). Four descriptors are sequence-based, computed from the Fv amino acid sequences: percent humanness (average variable-gene identity to the nearest human germline, from IgBLAST), length-normalized aggregation propensity (TANGO, at pH 7.4 and 150 mM salt), a chemical-liabilities penalty score (a weighted count of degradation-prone sequence motifs, with weights depending on framework-versus-CDR location), and intrinsic immunogenicity potential (the fraction of residues within promiscuous predicted MHC class II epitopes, from NetMHCIIpan across representative HLA-II alleles). The remaining five descriptors are structure-based, computed from homology models of each Fv built with the Antibody Modeler in MOE (protonated and energy-minimized): the surface area buried between the VH and VL domains (BSA_VH-VL_, a proxy for conformational stability), the structure-based isoelectric point (pI_Fv,3D_), hydrophobic imbalance (HI, the asymmetry in surface hydrophobic-residue distribution), the ratio of electrostatic dipole to hydrophobic moment (RM), and the ratio of charged to hydrophobic patch surface area (RP). All descriptors are computed at the physiological condition (pH 7.4, 150 mM ionic strength). Each raw descriptor is converted to a Z-score against the marketed-antibody reference distribution, with Z-scores capped at ±3 to limit outlier influence, and signed according to whether higher, lower, or nearer-to-mean values are favorable for developability. The four sequence-based and five structure-based transformed Z-scores are summed into a sequence score and a structure score, respectively, and these are added to yield a single total medicine-likeness score; the profile uses equal weighting across all nine descriptors. Antibodies with high sequence and structure scores are expected to show favorable intrinsic developability profiles. The score is a physicochemical enrichment heuristic and is not intended as a validated prospective predictor of experimental outcomes.

### 4.4 Experimental characterization

Expression, purification, and biophysical characterization were attempted for 255 antibodies in IgG1 format: 122 derived from variable regions of marketed therapeutics (as of February 2024), 93 GAN-generated, and 40 VAE-generated. Of these, 254 yielded complete biophysical records (one GAN candidate failed expression). The GAN and VAE generated antibodies were sampled from our refined models trained on highly medicine-like germline sequences. Candidates were produced as full-length human IgG1 monoclonal antibodies (mAbs) in CHO cells and, where expression succeeded, purified via Protein A affinity chromatography and size-exclusion chromatography and characterized across nine standardized biophysical assays. Assays included measurements of expression yield, aggregation propensity, thermal stability (Tm and Tagg), hydrophobicity (HIC retention time), poly-reactivity (DNA and insulin ELISA), and self-association (AC-SINS). Full experimental methods, assay conditions, and detailed results are provided elsewhere (Liu et al., in preparation).

### 4.5 Correlation analysis

To assess the relationship between computed medicine-likeness descriptors (sequence score, structure score, and total score) and experimental measurements, we calculated Pearson correlation coefficients between each pair of simulated and experimental properties. VAE-generated antibodies were excluded from this analysis due to their poor overall experimental performance. To quantify uncertainty in these correlation estimates, we employed bootstrap resampling with 10,000 iterations. For each iteration, we randomly sampled n antibodies with replacement (where n = 226, comprising all non-VAE antibodies with complete descriptor and assay data: 90 GAN-generated, 109 marketed, and 27 additional reference antibodies comprising 14 engineered point-mutant variants, 7 clinical-stage benchmark antibodies, 5 withdrawn therapeutics, and the NISTmAb reference standard), computed the Pearson correlation coefficient, and stored the result. The 95% confidence intervals were derived from the 2.5th and 97.5^th^ percentiles of the resulting bootstrap distribution. Because a subset of the reference antibodies are engineered variants of shared parent sequences, these correlation estimates should be read as descriptive rather than as inference over fully independent observations. Results are displayed in Fig. 4 (panel A) as a sparkline visualization where each cell shows the correlation coefficient as a central point with 95% confidence interval bounds represented by horizontal bars and a vertical reference line at zero.

### 4.6 Predictive modelling

#### Data Preparation and Feature Engineering

Antibody sequence data were collected for both marketed and GAN-generated antibodies. Each antibody was characterized using a set of computational molecular descriptors derived from the sequences: total medicine-like score, buried surface area (BSA) metrics, protein isoelectric point, dipole-hydrophobic ratio, patch ratio, hydrophobic index (HI), humanness scores, chemical liabilities, aggregation propensity, and immunogenicity risk scores. These computational descriptors served as input features for predictive modeling to assess whether computed properties could predict experimental outcomes.

#### Target Variables and Binning

Five experimental assay outcomes were selected as target variables: HIC-HPLC retention time, AC-SINS Score Δ*λ*_max_, DSF TM1, SLS Tagg, and protein yield. For classification tasks, continuous targets were discretized using three binning strategies: binary (median split), tertile (33rd and 67th percentiles), and quartile (25th, 50th, and 75th percentiles).

#### Sequence Similarity-Blocked Cross-Validation

To prevent data leakage from highly similar antibody sequences, a sequence similarity-based cross-validation approach was implemented. Pairwise sequence similarity was computed for all antibodies using combined Hamming distance across heavy and light chains. Heavy and light chain similarities were calculated as 1 minus the normalized Hamming distance, and combined with equal weighting. Sequences were grouped into clusters using connected components of a similarity graph, with thresholds of 0.8, 0.9, and 1.0. Train/test splits were constructed by randomly assigning entire clusters to either training or testing sets, ensuring that no test sequence exceeded the similarity threshold with any training sequence.

#### Model Training and Evaluation

Regression models were built using Ridge regression with standardized computational descriptors as features. Classification models used Random Forest classifiers with 100 estimators and a maximum depth of 10. Model performance was evaluated using *R*^2^, root mean squared error (RMSE), and mean absolute error (MAE) for regression, and macro-averaged F1 score, accuracy, and confusion matrices for classification. RMSE values were normalized by the standard deviation of each target variable. For each similarity threshold, three random seeds (42, 123, 456) were used to generate independent train/test splits.

#### Model Validation

For each split, the maximum similarity between test and training sequences was computed to verify adherence to the specified similarity constraints. All modeling and evaluation steps were repeated for each target variable and binning strategy to assess the predictive capacity of computational descriptors for experimental antibody properties.

## 5 Author contributions

Sandeep Kumar initiated the work and led the overall effort. Arkadij Kummer, Farbod Mahmoudinobar, Wendi Liu, J. Wade Davis, Eric J. Ma, and Sandeep Kumar contributed to the study and manuscript. All authors reviewed and approved the manuscript.

## Notes

### Competing Interest Statement

J. Wade Davis and Eric J. Ma are employees of Moderna, Inc. Farbod Mahmoudinobar was an employee of Moderna, Inc. during the conduct of this work. The authors declare no other competing interests.

### Funding

- Moderna, Inc.

## References

[1] Bauer, J., Rajagopal, N., Gupta, P., Gupta, P., Nixon, A.E., Kumar, S.: How can we discover developable antibody-based biotherapeutics? Frontiers in Molecular Biosciences 10 (2023). Accessed 2023-09-19

[2] Rajagopal, N., Choudhary, U., Tsang, K., Martin, K.P., Karadag, M., Chen, H.- T., Kwon, N.-Y., Mozdzierz, J., Horspool, A.M., Li, L., Tessier, P.M., Marlow, M.S., Nixon, A.E., Kumar, S.: Deep learning-based design and experimental validation of a medicine-like human antibody library. Briefings in Bioinformatics 26(1), 023 (2025) 10.1093/bib/bbaf023. Accessed 2025-11-10

[3] Giessel, A., Dousis, A., Ravichandran, K., Smith, K., Sur, S., McFadyen, I., Zheng, W., Licht, S.: Therapeutic enzyme engineering using a generative neural network. Scientific Reports 12(1), 1536 (2022) 10.1038/s41598-022-05195-x. Publisher: Nature Publishing Group. Accessed 2025-11-10

[4] Repecka, D., Jauniskis, V., Karpus, L., Rembeza, E., Rokaitis, I., Zrimec, J., Poviloniene, S., Laurynenas, A., Viknander, S., Abuajwa, W., Savolainen, O., Meskys, R., Engqvist, M.K.M., Zelezniak, A.: Expanding functional protein sequence spaces using generative adversarial networks. Nature Machine Intelligence 3(4), 324–333 (2021) 10.1038/s42256-021-00310-5. Publisher: Nature Publishing Group. Accessed 2025-11-10

[5] Kenlay, H., Dreyer, F.A., Kovaltsuk, A., Miketa, D., Pires, D., Deane, C.M.: Large scale paired antibody language models. PLOS Computational Biology 20(12), 1012646 (2024) 10.1371/journal.pcbi.1012646. Models: IgBert, IgT5. Preprint arXiv:2403.17889.

[6] Wang, Y., Bhattacharya, T., Jiang, Y., Qin, X., Wang, Y., Liu, Y., Saykin, A.J., Chen, L.: A novel deep learning method for predictive modeling of microbiome data. Briefings in Bioinformatics 22(3) (2021) 10.1093/bib/bbaa073. Accessed 2021-09-21

[7] Rajagopal, N., Choudhary, U., Tsang, K., Martin, K.P., Karadag, M., Chen, H.- T., Kwon, N.-Y., Mozdzierz, J., Horspool, A.M., Li, L., Tessier, P.M., Marlow, M.S., Nixon, A.E., Kumar, S.: Deep learning-based design and experimental validation of a medicine-like human antibody library. Briefings in Bioinformatics 26(1), 023 (2025) 10.1093/bib/bbaf023. Accessed 2025-11-10

[8] Wasdin, P.T., Johnson, N.V., Janke, A.K., Held, S., Marinov, T.M., Jordaan, G., Gillespie, R.A., Vandenabeele, L., Pantouli, F., Powers, O.C., Vukovich, M.J., Holt, C.M., Kim, J., Hansman, G., Logue, J., Chu, H.Y., Andrews, S.F., Kanekiyo, M., Sautto, G.A., Ross, T.M., Sheward, D.J., McLellan, J.S., Abu-Shmais, A.A., Georgiev, I.S.: Generation of antigen-specific paired-chain antibodies using large language models. Cell 188(25), 7206–722116 (2025) 10.1016/j.cell.2025.10.006. Model: MAGE. PMID 41192421; PMC12684077.

[9] Zhu, Y., Ma, J., Yin, M., Wu, J., Tang, L., Zhang, Z., Li, Q., Feng, S., Liu, H., Qin, T., Yan, J., Hsieh, C.-Y., Hou, T.: Ophiuchus-Ab: A Versatile Generative Foundation Model for Advanced Antibody-Based Immunotherapy. bioRxiv. bioRxiv 2026.02.02.703197; masked diffusion language model on paired chains, in silico only. (2026). 10.64898/2026.02.02.703197. https://www.biorxiv.org/content/10.64898/2026.02.02.703197v1

[10] Shuai, R.W., Ruffolo, J.A., Gray, J.J.: IgLM: Infilling language modeling for antibody sequence design. Cell Systems 14(11), 979–9894 (2023) 10.1016/j.cels.2023.10.001. Publisher: Elsevier. Accessed 2025-11-10

[11] Shanehsazzadeh, A., McPartlon, M., Kasun, G., Steiger, A.K., Sutton, J.M., Yassine, E., McCloskey, C., Haile, R., Shuai, R., Alverio, J., Rakocevic, G., Levine, S., Cejovic, J., Gutierrez, J.M., Morehead, A., Dubrovskyi, O., Chung, C., Luton, B.K., Diaz, N., Kohnert, C., Consbruck, R., Carter, H., LaCombe, C., Bist, I., Vilaychack, P., Anderson, Z., Xiu, L., Bringas, P., Alarcon, K., Knight, B., Radach, M., Bateman, K., Kopec-Belliveau, G., Chapman, D., Bennett, J., Ventura, A.B., Canales, G.M., Gowda, M., Jackson, K.A., Caguiat, R., Brown, A., Silva, D.G.d., Guo, Z., Abdulhaqq, S., Klug, L.R., Gander, M., Yapici, E., Meier, J., Bachas, S.: Unlocking de novo antibody design with generative artificial intelligence. bioRxiv. Pages: 2023.01.08.523187 Section: New Results (2024). 10.1101/2023.01.08.523187. https://www.biorxiv.org/content/10.1101/2023.01.08.523187v4 Accessed 2025-11-10

[12] Raybould, M.I.J., Deane, C.M.: The Therapeutic Antibody Profiler for Computational Developability Assessment. Methods in Molecular Biology (Clifton, N.J.) 2313, 115–125 (2022) 10.1007/978-1-0716-1450-15

[13] Armstrong, G.B., Shah, V., Sanches, P., Patel, M., Casey, R., Jamieson, C., Burley, G.A., Lewis, W., Rattray, Z.: A framework for the biophysical screening of antibody mutations targeting solvent-accessible hydrophobic and electrostatic patches for enhanced viscosity profiles. Computational and Structural Biotechnology Journal 23, 2345–2357 (2024) 10.1016/j.csbj.2024.05.041

[14] FreyLab, et al.: Loop: Large-scale antibody discovery via computational design and experimental validation. Nature Biotechnology (2025). In preparation

[15] Arsiwala, A., Bhatt, R., Yang, Y., Cadena, P.Q., Anderson, K.C., Ao, X., Niekerk, L.v., Rosenbaum, A., Bhatt, A., Smith, A., Grippo, L., Cao, X., Cohen, R., Patel, J., Allen, O., Faraj, A., Nandy, A., Hocking, J., Tural, B., Salvador, S., Jacobowitz, J., Schaven, K., Sherman, M., Shah, S., Tessier, P.M., Borhani, D.: A high-throughput platform for biophysical antibody developability assessment to enable AI/ML model training. bioRxiv. Pages: 2025.05.01.651684 Section: New Results (2025). 10.1101/2025.05.01.651684. https://www.biorxiv.org/content/10.1101/2025.05.01.651684v1 Accessed 2025-11-10

[16] Quality by Design for Preclinical In Vitro Assay Development - Jones – 2025 - Pharmaceutical Statistics - Wiley Online Library. https://onlinelibrary.wiley.com/doi/full/10.1002/pst.2430 Accessed 2025-08-12

[17] Arjovsky, M., Chintala, S., Bottou, L.: Wasserstein GAN. arXiv. arXiv:1701.07875 [stat] (2017). 10.48550/arXiv.1701.07875. http://arxiv.org/abs/1701.07875 Accessed 2025-11-10

[18] Gulrajani, I., Ahmed, F., Arjovsky, M., Dumoulin, V., Courville, A.: Improved Training of Wasserstein GANs. arXiv. arXiv:1704.00028 [cs] (2017). 10.48550/arXiv.1704.00028. http://arxiv.org/abs/1704.00028 Accessed 2025-11-10

[19] Seidler, C.A., Spanke, V.A., Gamper, J., Bujotzek, A., Georges, G., Liedl, K.R.: Data-driven analyses of human antibody variable domain germlines: pairings, sequences and structural features. mAbs 17(1), 2507950 (2025) 10.1080/19420862.2025.2507950. PMID 40413729; PMC12118439. Analyzes 1,954,079 paired OAS sequences.

[20] Dunbar, J., Deane, C.M.: ANARCI: antigen receptor numbering and receptor classification. Bioinformatics 32(2), 298–300 (2016) 10.1093/bioinformatics/btv552. Accessed 2025-11-10

[21] Briney, B., Inderbitzin, A., Joyce, C., Burton, D.R.: Commonality despite exceptional diversity in the baseline human antibody repertoire. Nature 566, 393–397 (2019) 10.1038/s41586-019-0879-y

[22] Mahmoudinobar, F., Meng, G., Davis, J.W., Kumar, S.: An intrinsic sequence-structural profile for mRNA-delivered therapeutic antibodies. Briefings in Bioinformatics 27(3), 284 (2026) 10.1093/bib/bbag284

[23] Sanders, J., Giancardo, L., Guo, L., Zhao, Y., Sonmez, K., Cheng, N., Yilmaz, M.: Conditional generation of antibody sequences with classifier-guided germline-absorbing discrete diffusion. arXiv. arXiv:2605.06720; model: MochiDiff. (2026). 10.48550/arXiv.2605.06720. https://arxiv.org/abs/2605.06720

[24] Ma, E.J., Kummer, A.: Principled Decision-Making Workflow with Hierarchical Bayesian Models of High-Throughput Dose-Response Measurements. Entropy 23(6), 727 (2021) 10.3390/e23060727. Number: 6 Publisher: Multidisciplinary Digital Publishing Institute. Accessed 2025-01-24

[25] Kohout, P., Vasina, M., Majerova, M., Novakova, V., Damborsky, J., Bednar, D., Marek, M., Prokop, Z., Mazurenko, S.: Engineering Dehalogenase Enzymes Using Variational Autoencoder-Generated Latent Spaces and Microfluidics. JACS Au 5(2), 838–850 (2025) 10.1021/jacsau.4c01101. Publisher: American Chemical Society. Accessed 2025-11-10

[26] Friedensohn, S., Neumeier, D., Khan, T.A., Csepregi, L., Parola, C., Vries, A.R.G.d., Erlach, L., Mason, D.M., Reddy, S.T.: Convergent selection in antibody repertoires is revealed by deep learning. bioRxiv. Pages: 2020.02.25.965673 Section: New Results (2020). 10.1101/2020.02.25.965673. https://www.biorxiv.org/content/10.1101/2020.02.25.965673v1 Accessed 2025-11-10

[27] Li, B., Luo, S., Wang, W., Xu, J., Liu, D., Shameem, M., Mattila, J., Franklin, M.C., Hawkins, P.G., Atwal, G.S.: Propermab: an integrative framework for in silico prediction of antibody developability using machine learning. mAbs 17(1), 2474521 (2025) 10.1080/19420862.2025.2474521

[28] Zou, W., Deng, J., Shen, Y., Zhang, T., Bing, Z., Yuan, L., Huang, C., Liu, J., Li, X.L.: Ab-panda: An ai-generated antibody structure-based tool for developability prediction. Biotechnology and Bioengineering 123(2), 287–297 (2026) 10.1002/bit.70086. Epub 2025 Nov 1. PMID 41174971. AlphaFold2-based developability prediction.

[29] Crouzet, S.J.: Biologically-Grounded Multi-Encoder Architectures as Developability Oracles for Antibody Design. arXiv. arXiv:2604.09369; model: CrossAbSense. (2026). 10.48550/arXiv.2604.09369. https://arxiv.org/abs/2604.09369

[30] Giancardo, L., Yilmaz, M., Lee, E., Ren, K., Zhao, Y., Trang, G., Sonmez, K., Guo, L., Cheng, N.: Context-aware Multi-Property Antibody Predictor: a Novel Framework Integrating Text and Protein Language Models. bioRxiv. bioRxiv 2026.01.07.698270; properties are computationally predicted, not experimentally measured. (2026). 10.64898/2026.01.07.698270. https://www.biorxiv.org/content/10.64898/2026.01.07.698270v1

[31] Zhao, S., Moller, J., Quintero-Cadena, P., Niekerk, L.: Guided Generation for Developable Antibodies. arXiv. arXiv:2507.02670; ICML 2025 GenBio Workshop. SVDD-guided generation. (2025). 10.48550/arXiv.2507.02670. https://arxiv.org/abs/2507.02670

[32] Shen, L., Feng, H., Qiu, Y., Wei, G.-W.: SVSBI: sequence-based virtual screening of biomolecular interactions. Communications Biology 6(1), 536 (2023) 10.1038/s42003-023-04866-3. Publisher: Nature Publishing Group. Accessed 2025-11-10

[33] Olsen, T.H., Moal, I.H., Deane, C.M.: AbLang: an antibody language model for completing antibody sequences. Bioinformatics Advances 2(1), 046 (2022) 10.1093/bioadv/vbac046. Accessed 2025-11-10

[34] Ruffolo, J.A., Gray, J.J., Sulam, J.: Deciphering antibody affinity maturation with language models and weakly supervised learning. arXiv. arXiv:2112.07782; model: AntiBERTy. (2021). 10.48550/arXiv.2112.07782. https://arxiv.org/abs/2112.07782

[35] Lin, Z., Akin, H., Rao, R., Hie, B., Zhu, Z., Lu, W., Smetanin, N., Verkuil, R., Kabeli, O., Shmueli, Y., Santos Costa, A., Fazel-Zarandi, M., Sercu, T., Candido, S., Rives, A.: Evolutionary-scale prediction of atomic-level protein structure with a language model. Science 379(6637), 1123–1130 (2023) 10.1126/science.ade2574. Model: ESM-2.

[36] Nijkamp, E., Ruffolo, J.A., Weinstein, E.N., Naik, N., Madani, A.: Progen2: Exploring the boundaries of protein language models. Cell Systems 14(11), 968–978 (2023) 10.1016/j.cels.2023.10.002

[37] Luo, S., Su, Y., Peng, X., Wang, S., Peng, J., Ma, J.: Antigen-specific antibody design and optimization with diffusion-based generative models for protein structures. In: Advances in Neural Information Processing Systems, vol. 35, pp. 9754–9767 (2022). 10.1101/2022.07.10.499510. Model: DiffAb.NeurIPS 35. 10.1101/2022.07.10.499510

[38] Martinkus, K., Ludwiczak, J., Cho, K., Liang, W.-C., Lafrance-Vanasse, J., Hotzel, I., Rajpal, A., Wu, Y., Bonneau, R., Gligorijevic, V., Loukas, A.: Abdiffuser: Full-atom generation of in-vitro functioning antibodies. In: Advances in Neural Information Processing Systems, vol. 36, pp. 40729–40759 (2023). 10.48550/arXiv.2308.05027. Model: AbDiffuser. NeurIPS 36; arXiv:2308.05027. https://arxiv.org/abs/2308.05027

[39] Cutting, D., Dreyer, F.A., Errington, D., Schneider, C., Deane, C.M.: De novo antibody design with SE(3) diffusion. arXiv. arXiv:2405.07622 [q-bio] (2024). 10.48550/arXiv.2405.07622. http://arxiv.org/abs/2405. 07622 Accessed 2025-11-10

[40] Dauparas, J., Anishchenko, I., Bennett, N., Bai, H., Ragotte, R.J., Milles, L.F., Wicky, B.I.M., Courbet, A., Haas, R.J., Bethel, N., Leung, P.J.Y., Huddy, T.F., Pellock, S., Tischer, D., Chan, F., Koepnick, B., Nguyen, H., Kang, A., Sankaran, B., Bera, A.K., King, N.P., Baker, D.: Robust deep learning-based protein sequence design using proteinmpnn. Science 378(6615), 49–56 (2022) 10.1126/science.add2187. Model: ProteinMPNN.

[41] Høie, M.H., Hummer, A., Olsen, T.H., Aguilar-Sanjuan, B., Nielsen, M., Deane, C.M.: AntiFold: Improved antibody structure-based design using inverse folding. arXiv. arXiv:2405.03370 [q-bio] (2024). 10.48550/arXiv.2405.03370. http://arxiv.org/abs/2405.03370 Accessed 2025-11-10

[42] Chai Discovery Team: Zero-shot antibody design in a 24-well plate. bioRxiv. bioRxiv 2025.07.05.663018, posted 2025-07-06; not peer reviewed. 16% de novo antibody design hit rate across 52 targets, over 100-fold improvement over prior computational methods. (2025). 10.1101/2025.07.05.663018. https://chaiassets.com/chai-2/paper/technicalreport.pdf

[43] Nabla Bio, Biswas, S.: De novo design of epitope-specific antibodies against soluble and multipass membrane proteins with high specificity, developability, and function. bioRxiv. bioRxiv 2025.01.21.633066; JAM (Joint Atomic Modeling) generative antibody design system. (2025). 10.1101/2025.01.21.633066. https://www.biorxiv.org/content/10.1101/2025.01.21.633066v1

[44] Latent Labs Team, Kenlay, H., Pretorius, D., Crabbe, J., Bridgland, A., Schmon, S.M., Hilmkil, A., Vuckovic, J., Mathis, S., Matteson, T., Bartke-Croughan, R., Motmaen, A., Rombach, R., Vlachynska, M., Nelson, A.W.R., Yuan, D., Obika, A., Kohl, S.A.A.: Drug-like antibodies with low immunogenicity in human panels designed with Latent-X2. arXiv. arXiv:2512.20263. Model: Latent-X2. (2025). https://arxiv.org/abs/2512.20263

[45] Zhao, X., Tang, Y.-C., Singh, A., Cantu, V.J., An, K., Lee, J., Stogsdill, A.E., Hamdi, I.M., Ramesh, A.K., An, Z., Jiang, X., Kim, Y.: AbBiBench: A Benchmark for Antibody Binding Affinity Maturation and Design. arXiv. arXiv:2506.04235 (2025). 10.48550/arXiv.2506.04235. https://arxiv.org/abs/2506.04235

[46] Ahmed, M., Taj, N., Khan, I.U., Venkateswara, H., Patterson, M.: CHIMERA-Bench: A Benchmark Dataset for Epitope-Specific Antibody Design. arXiv. arXiv:2603.13431; GEM workshop, ICLR 2026. Structure-conditioned sequence-structure co-design. (2026). 10.48550/arXiv.2603.13431. https://arxiv.org/abs/2603.13431

[47] Johnson, S.R., Fu, X., Viknander, S., Goldin, C., Monaco, S., Zelezniak, A., Yang, K.K.: Computational scoring and experimental evaluation of enzymes generated by neural networks. Nature Biotechnology 43(3), 396–405 (2025) 10.1038/s41587-024-02214-2. Publisher: Nature Publishing Group. Accessed 2025-11-10

